# The oncogene cyclin D1 promotes bipolar spindle integrity under compressive force

**DOI:** 10.1101/2023.05.30.542893

**Authors:** Renaldo Sutanto, Lila Neahring, Andrea Serra Marques, Seda Kilinc, Andrei Goga, Sophie Dumont

## Abstract

The mitotic spindle is the bipolar, microtubule-based structure that segregates chromosomes at each cell division. Aberrant spindles are frequently observed in cancer cells, but how oncogenic transformation affects spindle mechanics and function, particularly in the mechanical context of solid tumors, remains poorly understood. Here, we constitutively overexpress the oncogene cyclin D1 in human MCF10A cells to probe its effects on spindle architecture and response to compressive force. We find that cyclin D1 overexpression increases the incidence of spindles with extra poles, centrioles, and chromosomes. However, it also protects spindle poles from fracturing under compressive force, a deleterious outcome linked to multipolar cell divisions. Our findings suggest that cyclin D1 overexpression may adapt cells to increased compressive stress, contributing to its prevalence in cancers such as breast cancer by allowing continued proliferation in mechanically challenging environments.

## Introduction

The spindle is the macromolecular machine that segregates chromosomes at each cell division. In mammalian cells, mitotic spindles are bipolar structures with one centrosome at each spindle pole. Errors in cell division are associated with genomic instability and disease, and aberrant spindles are hallmarks of cancer (Holland and Cleveland, 2009). Extra centrosomes (Lingle *et al*., 1998; Pihan *et al*., 1998; Chan, 2011), continuously evolving karyotypes known as chromosomal instability (Lengauer *et al*., 1997; Drews *et al*., 2022), and multipolar spindles are elevated in tumors across many tissues of origin and diverse cancer genotypes. Oncogenes can also induce defects in spindle assembly even in the absence of gross spindle abnormalities; for example, MYC overexpression prolongs mitosis and increases chromosome segregation errors (Rohrberg *et al*., 2020). Paradoxically, while such multipolar, clustered pseudo-bipolar, or otherwise aberrant spindles are generally adverse for mitotic outcomes (Ganem *et al*., 2009; Silkworth *et al*., 2009), they can promote tumorigenesis by increasing genetic diversity (Holland and Cleveland, 2009) and potentially other unknown mechanisms. How oncogenic transformation affects spindle assembly remains poorly understood.

Dividing cells in solid tumors are subject to dramatically different mechanical environments than their counterparts in healthy tissue (Kumar and Weaver, 2009; Plodinec *et al*., 2012; Nia *et al*., 2016). Spindle poles in dividing cultured cells often fracture under compressive force, leading to mitotic delays, multipolar anaphases, and subsequent cell death (Tse *et al*., 2012; Lancaster *et al*., 2013; Matthews *et al*., 2020; Cheng *et al*., 2023). Tumors have been shown to be confining microenvironments due to their increased cell density, elevated interstitial fluid pressure (Heldin *et al*., 2004), and increased extracellular matrix deposition and crosslinking (Levental *et al*., 2009), raising the question of how cells continue to divide under this high compressive stress. In breast tumors, compressive stress is high enough to deform and damage interphase nuclei (Nader *et al*., 2021), and nearby mitotic cells presumably experience similarly high forces that may interfere with mitotic rounding or spindle assembly. In multicellular tumor spheroid models, compressive stress reduces cell proliferation (Helmlinger *et al*., 1997; Cheng *et al*., 2009; Delarue *et al*., 2014; Taubenberger *et al*., 2019) and has been shown to disrupt bipolar spindle assembly in cells that continue to divide (Desmaison *et al*., 2013). Due to the challenges of making controlled mechanical perturbations at the cellular scale, little is known about whether and how the spindles of transformed cells mechanically differ from wild-type spindles as they adapt to the tumor environment.

Cyclin D1, overexpressed in 50-70% of breast cancers (Musgrove *et al*., 2011), is an oncogene with pleiotropic effects in the cell. Acute overexpression of cyclin D1 leads to spindle and karyotypic defects (Nelsen *et al*., 2005), and long-term overexpression is sufficient to drive breast cancer in mice (Wang *et al*., 1994). In addition to its canonical role in complex with CDK4/6 in controlling cell cycle progression at the G1/S transition, cyclin D1 may contribute to tumorigenesis through its roles in cytoskeletal remodeling and CDK-independent transcriptional programs (Musgrove *et al*., 2011). Many other oncogenes commonly dysregulated in breast cancer, such as Ras and ErbB2, are upstream of cyclin D1 (Lee *et al*., 2000; Yu *et al*., 2001; Desai *et al*., 2002), making cyclin D1 overexpression a good model to probe changes in spindle mechanics after oncogenic transformation.

Here, we compare control and cyclin D1-overexpressing breast epithelial cells to investigate their spindle architectures and responses to compressive stress. We find that cyclin D1 increases the proportion of spindles containing extra poles, chromosomes, and centrosomes. However, cyclin D1 overexpression also promotes bipolar spindle integrity during cell compression, preventing spindle pole fracture that results in multipolar cell divisions. We propose that cyclin D1 overexpression mechanically adapts cell division to the tumor context, potentially contributing to its prevalence in cancer despite the aberrant spindles it induces.

## Results and Discussion

### Constitutive cyclin D1 overexpression promotes aberrant spindle architectures

To determine the effects of cyclin D1 overexpression on spindle architecture, we compared MCF10A breast epithelial cell lines stably overexpressing cyclin D1 or a puromycin resistance gene as a control (Figure 1A) (Martins *et al*., 2015). The parental MCF10A cells are diploid and non-transformed, but are sensitive to transformation by a variety of oncogenes (Soule *et al*., 1990; Debnath *et al*., 2002; Martins *et al*., 2015). We confirmed overexpression of cyclin D1 by western blot (Figure 1B), and used immunofluorescence to quantify spindle pole, centriole, and kinetochore numbers by staining for α-tubulin, centrin, and CREST respectively (Figure 1C).

**Figure 1.**
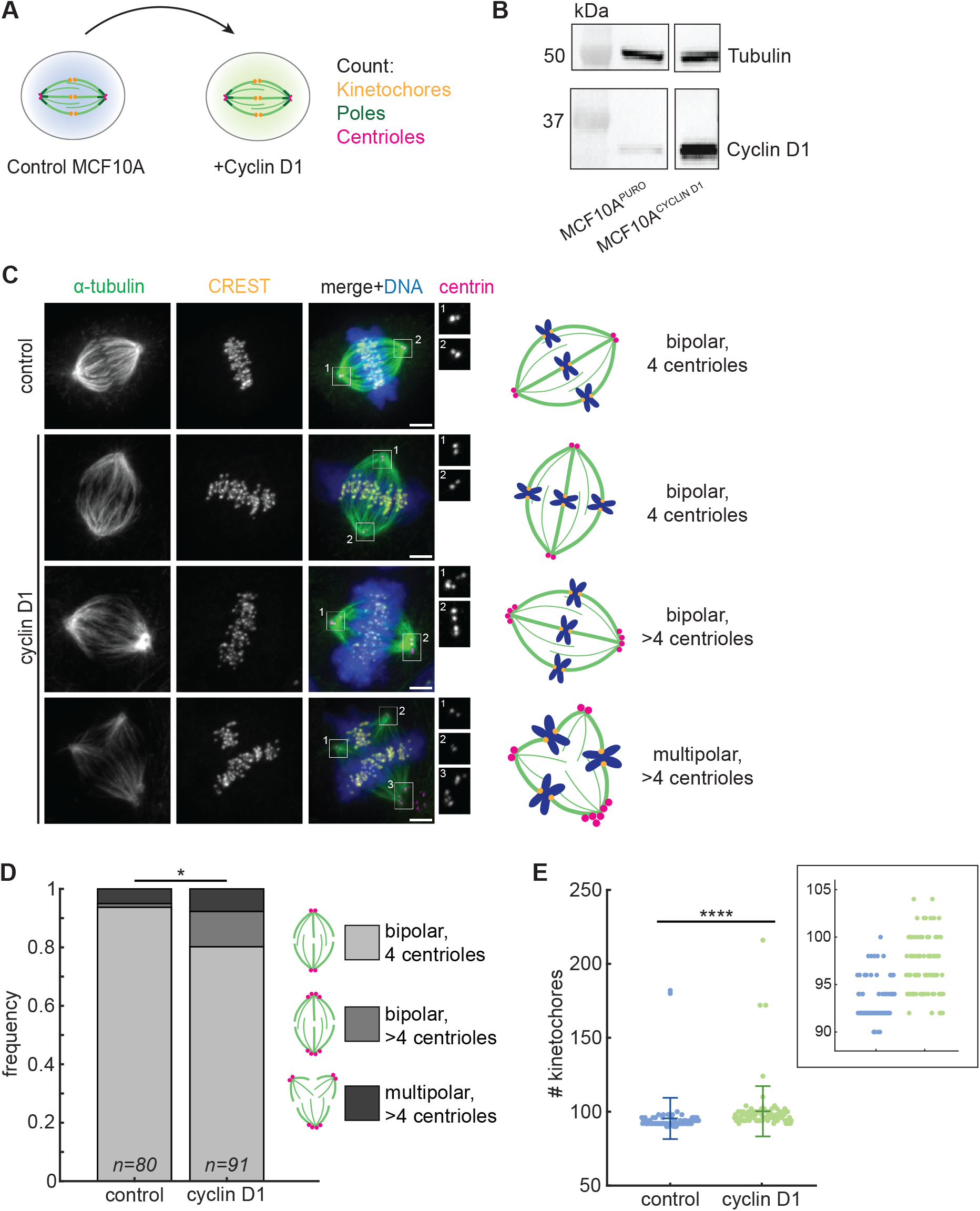
Cyclin D1 overexpression promotes aberrant spindle architectures. (A) Schematic diagram of assay. MCF10A cells stably expressing a puromycin resistance gene (control) or cyclin D1 with puromycin resistance were examined by immunofluorescence for changes to metaphase spindle pole, centriole, or kinetochore numbers. (B) Western blot of α-tubulin and cyclin D1 levels in MCF10A^PURO^ (control) and MCF10A^CYCLIN^ ^D1^ cell lines. All images are from the same blot, with intervening lanes removed. (C) Representative confocal immunofluorescence images (maximum intensity projections) of spindles stained for α-tubulin (green), CREST (yellow), centrin (magenta), and Hoechst (blue), with spindle phenotypes cartooned (right). Magnifications of the centrioles at each spindle pole are shown at right. Scale bars = 3 µm. (D) Frequency of the three observed metaphase spindle phenotypes in each MCF10A cell line. The distribution of phenotypes differs between cyclin D1 and control cells (*p = 0.010, Fisher’s exact test), with the cyclin D1 line enriched in cells with supernumerary centrioles. (E) Number of kinetochores per spindle. Metaphase spindles in the cyclin D1 cell line had significantly more kinetochores (representing the number of chromatids) than the control line (****p = 2.32×10^-14^, Mann-Whitney U test). Lines indicate mean ± standard deviation. Inset shows a smaller range of kinetochore numbers. For D and E, n = 80 control spindles and 91 cyclin D1 spindles, each pooled from 3 independent experiments.

While most control spindles (94%) had two centrioles at each of two spindle poles, supernumerary centrioles were more common in the cyclin D1-expressing cells (20% of cells; Figure 1D). These centrioles were either associated with multipolar spindles or clustered into pseudo-bipolar spindles, a known mechanism by which cancer cells adapt to extra centrosomes in order to avoid multipolar divisions (Ring *et al*., 1982; Quintyne *et al*., 2005; Kwon *et al*., 2008).

To gain insight into cyclin D1’s effect on genomic integrity, we next counted the kinetochores in each spindle. Several mechanisms, including the clustering of extra centrosomes (Ganem *et al*., 2009; Silkworth *et al*., 2009) and reduced kinetochore-microtubule dynamics (Bakhoum *et al*., 2009), have been shown to give rise to aneuploidy and chromosomal instability in cancer cells, while cytokinesis failure leads to larger-scale genomic duplications. Cyclin D1 overexpression was associated with a broader range of chromosome numbers than in controls, with only a small number of cells containing a near-doubling of chromosome number, indicating that it induces aneuploidy (Figure 1E). In summary, constitutive overexpression of the oncogene cyclin D1 leads to an increased incidence of spindles with extra poles, centrioles, and chromosomes, even when cells are allowed to adapt to elevated cyclin D1 over many passages.

### Cyclin D1 overexpression promotes bipolar spindle integrity under compressive stress

Although cyclin D1 overexpression gave rise to higher rates of spindle defects (Figure 1), it is overexpressed in many tumors such as breast cancers where dividing cells are subject to increased compressive stress (Musgrove *et al*., 2011; Nia *et al*., 2016; Nader *et al*., 2021). To account for its high prevalence as an oncogene, we hypothesized that cyclin D1 overexpression may alter the spindle’s biophysical properties in a manner that is adaptive in the tumor environment. We compared the mechanical robustness of control versus cyclin D1-overexpressing spindles by compressing cells in PDMS-based microfluidic devices and performing live imaging (Figure 2A) (Le Berre *et al*., 2014). Cells were pre-treated with the proteasome inhibitor MG132 to prevent anaphase entry, allowing us to focus on the metaphase spindle’s response to compressive stress, and gradually compressed to a final height of 5 µm via a computer-controlled vacuum pump over 4 minutes. Compression was then sustained for an additional 70 minutes. This perturbation was reproducible from cell to cell, reducing spindle height from an average of 10.55 ± 1.54 µm to 4.72 ± 0.31 µm (mean ± standard deviation of all cells) (Figure 2, B and C). Spindles in control and cyclin D1-overexpressing cells had indistinguishable average heights prior to compression and were compressed to a similar final height (Figure 2C). Spindles also widened and elongated as compression was applied, consistent with previous work (Dumont and Mitchison, 2009; Guild *et al*., 2017; Neahring *et al*., 2021). Spindle lengths before compression were similar between the control and cyclin D1 cells, as were spindle lengths at 10 minutes post-compression onset, when spindle shape had stabilized (Figure 2D). Spindles were significantly wider in control cells vs. cyclin D1-overexpressing cells, both before and after compression, but the difference was slight (Figure 2E). Thus, our assay directly probes the spindle’s intrinsic ability to adapt to a confined geometry under cyclin D overexpression, rather than probing the cell’s ability to shield the spindle from shape changes under compression.

**Figure 2.**
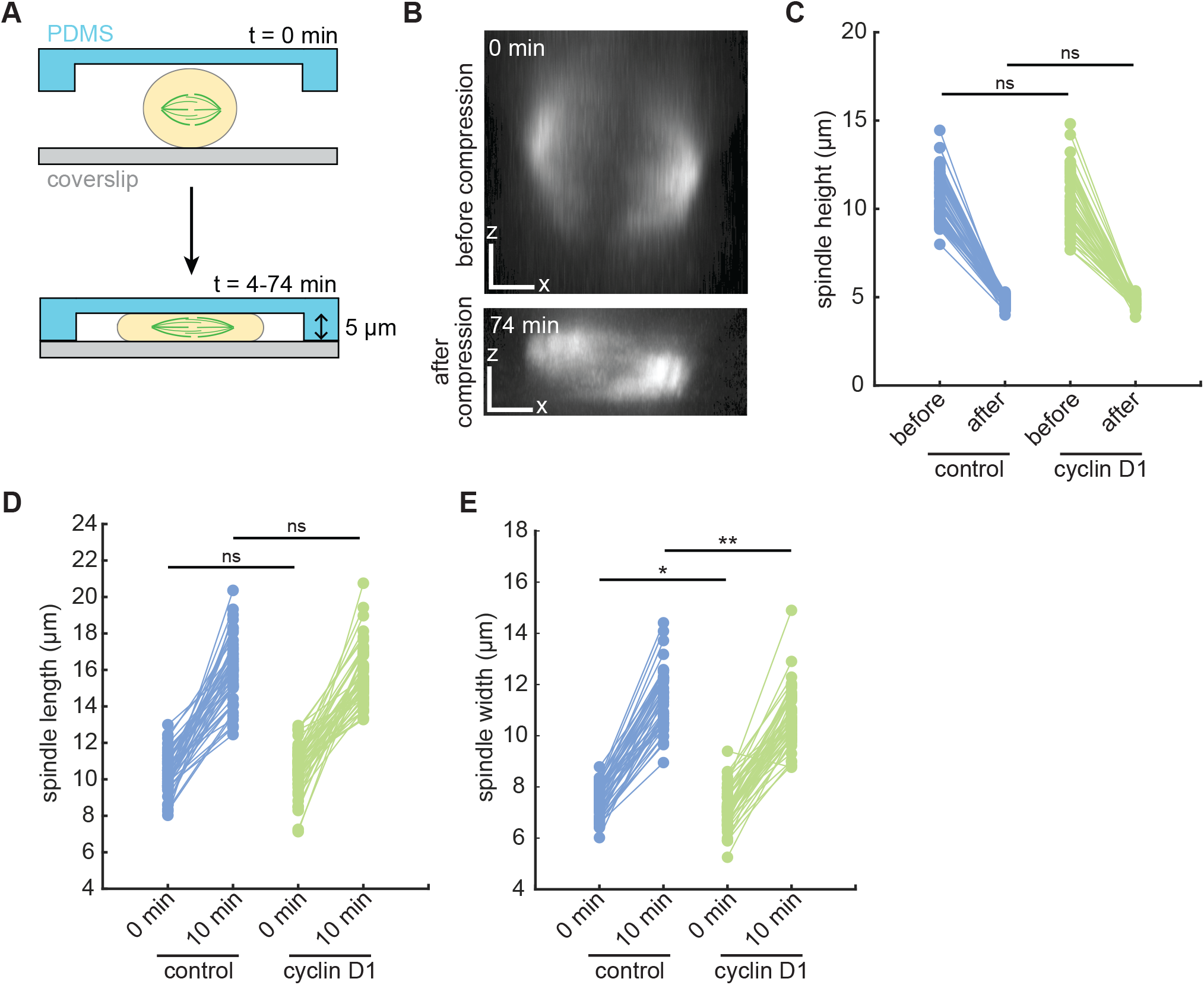
The cell compression assay is quantitatively reproducible. (A) Schematic diagram of cell compression assay using a microfluidic device. Cells were compressed to a height of 5 µm using computer-controlled negative pressure over 4 min, and compression was sustained for 70 additional minutes. Cells were live-imaged throughout to monitor changes in spindle architecture. (B) Side (XZ) views of a control spindle, labeled with SiR-tubulin, before and after (at 74 min) compression. X and Z scale bars = 3 µm. (C) Between the control and cyclin D1 cell lines, spindle heights did not significantly differ before compression, and spindles were compressed to a similar final height (measured at 74 min). ns, not significant. (D) Spindle lengths before and 10 minutes after compression onset (ns, not significant). (E) Spindle widths before and 10 minutes after compression onset (*p = 0.028, **p = 0.00066). For C-E, two-tailed two-sample t-tests were performed with n = 57 control and 53 cyclin D1 spindles (C) or n = 48 control and 52 cyclin D1 spindles (D and E). Spindles were excluded from length and width analysis if both poles were not in focus in the same z-plane.

During compression experiments, we monitored changes in spindle integrity in addition to changes in spindle shape. Control spindle poles fractured into multiple foci during the 74 minutes of compression 47.4% of the time, with kinetochore-fibers detaching and splaying laterally from the original spindle pole, and the metaphase plate becoming bent (Figure 3, A and B; Movie S1). Interestingly, bipolar spindles in the cyclin D1-overexpressing line fractured significantly less often, in just 20.8% of compressions (Figure 3B; Movie S2). Although these spindles experienced similar compression-induced deformations, most spindles maintained all kinetochore-fibers focused into the two original spindle poles, with the metaphase plate remaining as a straight line.

**Figure 3.**
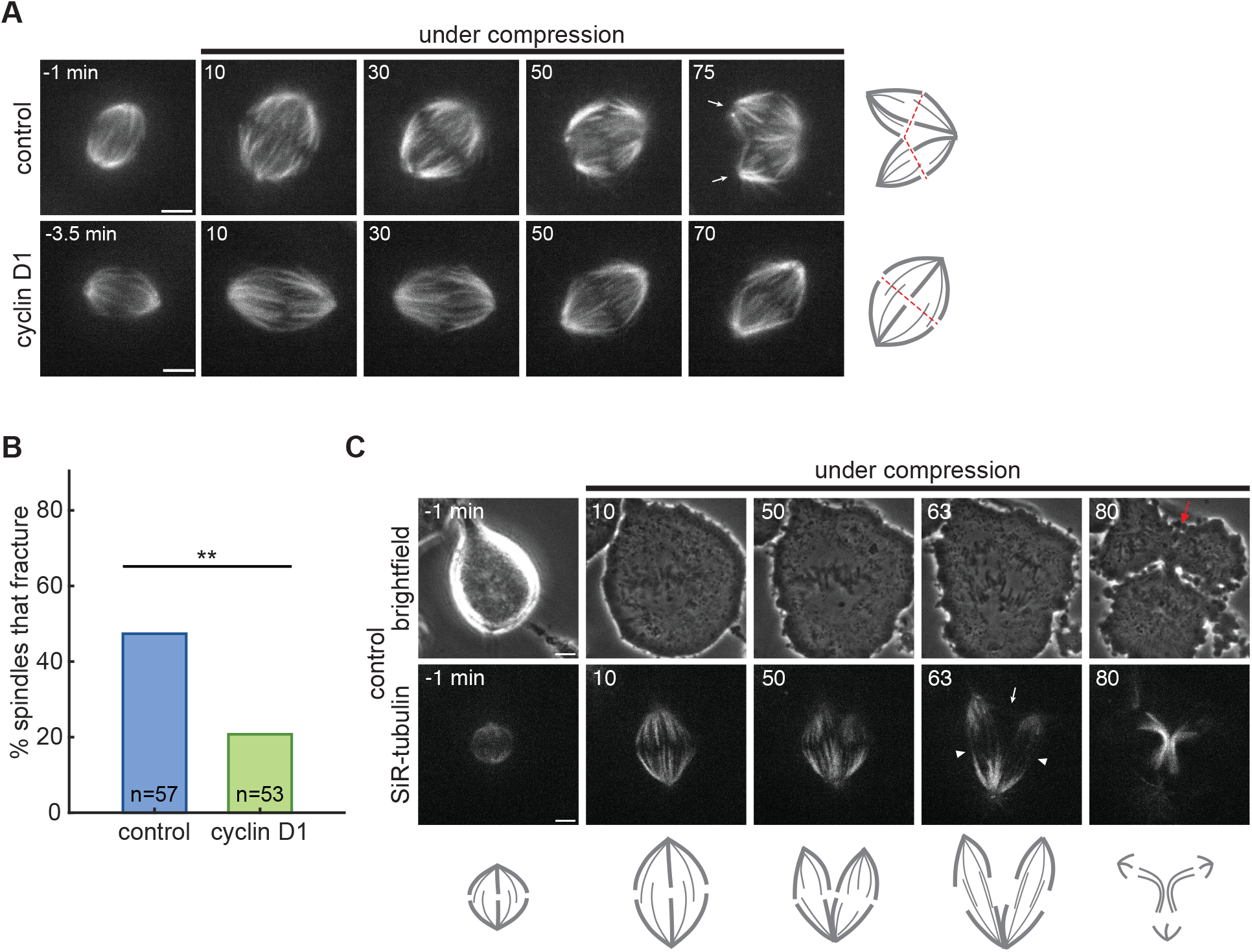
Cyclin D1 overexpression protects against spindle pole fracture during compression. (A) Confocal time-lapse images of control and cyclin D1-overexpressing cells undergoing compression, where the control spindle fractures around 60 min after compression onset (cartooned at right; dashed red lines represent the metaphase plate position). The fractured poles of the control spindle are indicated in the final frame by white arrows. Tubulin is labeled with SiR-tubulin. Scale bars = 5 µm; time stamps are in minutes. (B) Spindles in cyclin D1-overexpressing cells fractured less often than control spindles during the 74 minutes of compression (**p = 0.0048, Fisher’s exact test). n = 57 control and 53 cyclin D1 spindles. (C) Confocal time-lapse images of a control spindle undergoing compression to a height of 4 µm, without the addition of MG132. After the spindle fractures between the 10 and 50 min time points, the cell enters anaphase and segregates chromosomes into 3 masses (cartooned below). Interpolar microtubule bundles connect the original pole to each of the fractured poles (white arrowheads), while no interpolar bundles connect the two poles resulting from the fracture (white arrow). The cytokinetic furrow is disrupted between the two fractured poles by the final 80 min timepoint (red arrow). Scale bars = 5 µm; time stamps are in minutes.

To probe the consequences of spindle pole fracture, we imaged control cell compressions without the addition of MG132 to follow spindles into late mitosis.

Fractured spindles were still able to progress to anaphase, but they segregated chromosomes into three or more masses (consistent with a bent metaphase plate), depending on the number of new poles created by fracture (Figure 3C; Movie S3; note the example shown was compressed using 4 µm micropillars). Interestingly, poles that separated from each other as a result of fracture were directly connected by few or no microtubules (Figure 3C, white arrow), and cytokinetic furrowing between these fractured poles was disrupted (Figure 3C, red arrow). Our results suggest that compressive force on mitotic MCF10A cells often causes spindle poles to fracture, leading to abnormal chromosome segregation at anaphase, but that overexpression of the oncogene cyclin D1 is protective against spindle fracture.

### Conclusions

Many oncogenes induce aberrant spindle architectures, yet they also promote uncontrolled cell proliferation in tumorigenesis. One explanation for this apparent paradox is that the elevated rate of chromosome mis-segregation in these spindles accelerates genome evolution and gives some cells a selective advantage (Ben-David and Amon, 2020). Here, we describe another mechanism that could contribute to the proliferative advantage induced by some oncogenes. Overexpression of cyclin D1 increases the prevalence of mitotic cells containing extra poles, centrioles, and chromosomes (Figure 1), but also reduces the frequency of spindle fracture under compressive stress (Figures 2-3). Our assay was conducted in two-dimensional culture and with compressive force that may differ in magnitude and direction from that experienced by cells in vivo. Indeed, a recent study using HeLa cells found that while confinement-induced cell flattening led to increased pole fracturing, confining cells into elongated, narrow channels was protective against pole fracturing (Cheng *et al*., 2023). However, an increase in spindle multipolarity has also been observed in confined HCT116 colorectal cancer cell spheroids (Desmaison *et al*., 2013), suggesting that our assay mimics compressive forces that exist in a crowded three-dimensional environment.

Although the fractured spindles we followed into anaphase segregated chromosomes into more than two masses (Figure 3C), many of these mitoses presumably resolved into two daughter cells due to the lack of an anaphase central spindle competent to recruit the cytokinetic machinery between the newly separated poles. However, rapid nuclear envelope reformation at mitotic exit may prevent these multiple DNA masses from merging and lead to genomic instability or cell cycle arrest. Because cyclin D1 overexpression has a protective effect on bipolar spindle integrity under compressive force, we propose that it helps to prevent multipolar anaphases and may allow cells to continue proliferating under compressive stress in the tumor context. Intriguingly, the cyclin D1 interactors pRb, p27, and p21 have been shown to mediate a G1 arrest in cells subjected to compressive stress (Delarue *et al*., 2014; Nam *et al*., 2019; Taubenberger *et al*., 2019), suggesting that cyclin D1 levels may affect the likelihood both that cells will continue to divide under compression and that they will complete these divisions successfully.

This work poses the question of the mechanisms by which cyclin D1 overexpression protects against spindle fracture. Cyclin D1 could regulate other factors involved in the cell’s or the spindle’s response to compression through its kinase-dependent or transcriptional roles (Musgrove *et al*., 2011). Pharmacologically inhibiting CDK4/6, the partner kinases of cyclin D1, and testing whether spindles in cyclin D1- overexpressing cells are sensitized to compressive stress could help determine whether cyclin D1’s pole protective effect is kinase dependent. Any effects of CDK4/6 inhibitors on spindle mechanics may also be therapeutically relevant, because these inhibitors are widely used to treat metastatic estrogen receptor-positive breast cancer (Pernas *et al*., 2018).

Indirect consequences of cyclin D1 overexpression could also underlie the spindle pole protection we observe. Incomplete mitotic rounding has been shown to lead to pole fracturing (Lancaster *et al*., 2013), and oncogenic h-Ras^G12V^ has been shown to prevent pole fracture in MCF10A cells by enhancing mitotic rounding under stiff gels (Matthews *et al*., 2020). We propose that different mechanisms are at play in the protective effect we observe here, because spindles in cyclin D1-overexpressing cells underwent fewer fractures despite being compressed to the same flattened height as spindles in control cells (Figure 2C). Supernumerary centrioles could contribute to the protective effect of cyclin D1 by increasing the density of microtubules and/or pericentriolar material at poles (Lingle *et al*., 1998; Godinho *et al*., 2014; Cosenza *et al*., 2017). Indeed, the proportion of bipolar spindles containing extra centrioles was increased from 1.3% of controls to 13.1% in the cyclin D1 cell line (Figure 1D), and whether the pole-protective effect of cyclin D1 occurs specifically in cells with centriole amplification is an important question. Finally, other proteins that are differentially regulated during oncogenic transformation (but not specifically downstream of cyclin D1) could affect pole integrity. For example, TPX2 and chTOG, proteins required for spindle pole integrity, are commonly upregulated in cancer (Garrett *et al*., 2002; Gergely *et al*., 2003; Carter *et al*., 2006). Future work dissecting the mechanism(s) by which cyclin D1 promotes bipolar spindle integrity under compression will be important to predict how generalizable this phenomenon is likely to be among tumors with diverse driver oncogenes. More broadly, achieving this goal will require a better understanding of the physical and molecular basis of spindle mechanical integrity (Gatlin *et al*., 2010; Shimamoto *et al*., 2011; Suresh *et al*., 2020).

The biochemical hallmarks of cancer, including anti-apoptotic signaling, metabolic reprogramming, and cell cycle dysregulation, are well-established (Hanahan and Weinberg, 2011). By contrast, our knowledge of the biophysical hallmarks of cancer lags behind, and addressing this gap could reveal new insights into disease progression. Our application of controlled, cellular-scale force suggests that cyclin D1 overexpression may adapt dividing cells to the mechanical burdens of the tumor environment. Better understanding the biophysical adaptations of cancer cells could lead to new ways to selectively target these cells for therapeutic gain.

## Materials and methods

### Cell culture

MCF10A^PURO^ and MCF10A^CYCLIN^ ^D1^ cells were created in a previous study (Martins *et al*., 2015). Both cell lines were cultured at 37°C and 5% CO_2_, and maintained in DMEM/F12 (Invitrogen) supplemented with 5% horse serum (Gibco), 20 ng/ml epidermal growth factor, 10 µg/ml insulin, 0.5 µg/ml hydrocortisone, 100 ng/ml cholera toxin, 100 U/ml penicillin, and 100 U/ml streptomycin. For immunofluorescence experiments, cells were plated on 25 mm round #1.5 coverslips, coated with poly-L- lysine and 0.1% gelatin solution, two days prior to fixation. For compression experiments, cells were plated in 35 mm petri dishes containing 23 mm #1.5 poly-D-lysine-coated coverslips (World Precision Instruments) two days prior to imaging. Cells were plated to achieve a confluency of ∼40-50% at imaging, to allow space for cells to expand under compression.

### Western blotting

Cells in 6-well plates were lysed, and protein extracts were collected after centrifugation at 4°C for 30 min. Protein concentrations were measured using a Bradford assay, and equal concentrations of each sample were separated on a 4-12% Bis-Tris gel (Invitrogen) by SDS-PAGE and transferred to a nitrocellulose membrane. Membranes were blocked with 4% milk, incubated in primary antibodies overnight at 4°C, and incubated with HRP-conjugated secondary antibodies for 1 hour. Proteins were detected using SuperSignal West Pico or Femto chemiluminescent substrates (Thermo Fisher). The following primary antibodies were used: mouse anti-α-tubulin DM1α (1:5000, Sigma-Aldrich T6199) and rabbit anti-cyclin D1 SP4 (1:1000, Abcam ab16663). The following secondary antibodies were used at a 1:10,000 dilution: goat anti-mouse IgG-HRP (Santa Cruz Biotechnology sc-2005) and mouse anti-rabbit IgG-HRP (Santa Cruz Biotechnology sc-2357).

### Immunofluorescence

Cells were fixed in cold methanol for 2 minutes at -20°C. Cells were washed in TBST (0.05% Triton-X 100 in TBS) and blocked with 2% BSA in TBST. Primary and secondary antibodies were diluted in TBST + 2% BSA and incubated for one hour at room temperature (primary antibodies) or 50 minutes at room temperature (secondary antibodies). DNA was labeled with 1 µg/ml Hoechst 33342 prior to mounting on slides with ProLong Gold Antifade Mountant (Thermo Fisher P36934). The following primary antibodies were used: mouse anti-α-tubulin DM1α conjugated to Alexa Fluor 488 (1:100, Cell Signaling Technologies 8058S), mouse anti-centrin clone 20H5 (1:200, Sigma-Aldrich 04-1624), and human anti-centromere protein CREST antibody (1:25, Antibodies Incorporated 15-234). Normal mouse IgG (1:100, Santa Cruz Biotechnology sc-2025) was used as a block before incubating in pre-conjugated mouse anti-α-tubulin DM1α Alexa Fluor 488. The following secondary antibodies were used: goat anti-mouse conjugated to Alexa Fluor 488 and 568 (1:400, Invitrogen A11001 and A11004) and goat anti-human conjugated to Alexa Fluor 647 (1:400, Invitrogen A21445).

### Cell compression

Cell compressions were performed using a 1-well dynamic cell confiner with 5 µm PDMS micropillars, or 4 µm micropillars for the example shown in Figure 3C (4DCell). The device was attached to an AF1 Dual vacuum/pressure controller (Elveflow) and negative pressure was controlled using the Elveflow ESI software. Prior to imaging, a seal was established between the compression device and the dish of cells by applying a negative pressure of -10 mbar. At the start of imaging, a linear pressure ramp was applied from -10 to -150 mbar over a period of 4 minutes to lower the pillared coverslip onto the cells. Once the PDMS pillars contacted the dish, compression was maintained for 70 minutes. Z-stacks were acquired before each compression and after each timelapse acquisition to determine spindle height before and after each compression and quantitatively compare compression outcomes.

### Imaging

Live imaging experiments were conducted in a stage-top humidified incubation chamber (Tokai Hit WSKM) maintained at 37°C and 5% CO_2_. In compression experiments, microtubules were labeled with 100 nM SiR-tubulin (Cytoskeleton, Inc.) and 10 µM verapamil for 30-60 minutes prior to imaging. For all compression experiments shown except for the example in Figure 3C, the proteasome inhibitor MG132 was added to a final concentration of 10 µM 10 minutes prior to imaging to prevent anaphase entry during compressions. All live and immunofluorescence imaging was performed on an inverted spinning disk confocal (CSU-X1, Yokogawa Electric Corporation) microscope (Eclipse Ti-E, Nikon) with the following components: head dichroic Semrock Di01-T405/488/568/647; 405 nm (100 mW), 488 nm (150 mW), 561 nm (100 mW), and 642 nm (100 mW) diode lasers; ET455/50M, ET525/50M, ET600/50M, ET690/50M, and ET705/72M emission filters (Chroma Technology); and a Zyla 4.2 sCMOS camera (Andor Technology). Exposures of 50-200 ms were used for fluorescence. Images were acquired with a 100× 1.45 NA Ph3 oil objective using MetaMorph 7.7.8.0 (Molecular Devices).

### Data and statistical analysis

Immunofluorescence images show maximum intensity projections (Figure 1C) and time strip images show single spinning disk confocal Z-slices (Figure 3). All images were formatted for publication using FIJI (Schindelin *et al*., 2012). The brightness/contrast for each channel was scaled identically between different example cells for immunofluorescence images. The brightness/contrast for videos and time strips were scaled individually to account for variations in tubulin labeling efficiency.

Kinetochores were counted using the multi-point tool in FIJI. For compression experiments, spindle heights were measured from XZ views generated from z-stacks (see Figure 2B) in a vertical direction perpendicular to the coverslip. A fracture was defined as the development of a clear gap in tubulin intensity between a kinetochore-fiber minus-end and the main spindle pole within 74 minutes of compression onset (Figure 3). Fisher’s exact test was used to compare categorical datasets (Figures 1D and 3B); two-tailed two-sample t-tests were used to compare the numerical datasets in Figure 2, C-E based on the assumption that spindle heights, lengths, and widths are approximately normally distributed; and a Mann-Whitney U test was used to compare the numerical dataset in Figure 1E due to kinetochore number distributions that deviated from a normal distribution. Statistical tests were performed using the ttest2, ranksum, and fishertest functions in MATLAB R2022b, and the fisher.test function in R for the 2×3 comparison in Figure 1D. P-values are given in the figure legends.

## Acknowledgements

We thank Rachel Nakagawa and Julia Rohrberg for experimental advice, and other members of the Goga and Dumont labs for helpful discussions. This work was supported by an NSF Graduate Research Fellowship, Fannie and John Hertz Foundation Fellowship, and the UCSF Discovery Fellows Program (L.N.); NIHR35GM136420, NSF 1548297 Center for Cellular Construction, Chan Zuckerberg Biohub, and the UCSF Byers Award (S.D.); the Mark Foundation and Atwater Family Foundation (A.G.); and NIH 5T32CA108462 (S.K.).

## Figure Legends

**Movie S1**. A control cell before and after compression. Time stamps are in min:sec, scale bars are 5 µm, and video playback is 30 frames per second. See also Figure 3A.

**Movie S2**. A cyclin D1-overexpressing cell before and after compression. Time stamps are in min:sec, scale bars are 5 µm, and video playback is 30 frames per second. See also Figure 3A.

**Movie S3**. A control cell before and after compression to a height of 4 µm, without the addition of MG132. Time stamps are in min:sec, scale bars are 5 µm, and video playback is 30 frames per second. See also Figure 3C.

## Notes

### Competing Interest Statement

The authors have declared no competing interest.

